# Temporal saliency for motion direction may arise from visual stimulus-specific adaptation in avian midbrain inhibitory nucleus

**DOI:** 10.1101/2021.11.07.467641

**Authors:** Jiangtao Wang, Shuman Huang, Zhizhong Wang, Songwei Wang, Li Shi

## Abstract

Food and predators are the most noteworthy objects for the basic survival of wild animals. In nature, both of these are often rare or deviant in both spatial and temporal domains and would soon attract an animal’s attention. Although stimulus-specific adaptation (SSA) is considered to be one neural basis of salient sound detection in the temporal domain, related research on visual SSA is lacking. The avian nucleus isthmi pars magnocellularis (Imc), which plays an extremely important role in the selective attention network, is one of the best models for investigating the neural correlate of visual stimulus-specific adaptation (SSA) and detection of salient stimulus in the temporal domain. Here, we used a constant order paradigm to test the existence of SSA in the pigeon’s Imc. We found that the strength of response of Imc neurons significantly decreased after repetitive motion stimuli, but recovered when the motion was switched to a novel direction, leading to the saliency detection of the novel motion direction. These results suggest that the inhibitory nucleus Imc shows visual SSA to motion direction, allowing the Imc to implement temporal saliency mapping and to determine the spatial-temporal saliency of the current stimulus. This also implies that pigeons may detect novel spatial-temporal stimuli during the early stage of sensory processing.

## Introduction

In the natural world, the capacity to respond rapidly to surprise, and especially to dangerous stimuli, is vitally important for an animal’s survival (Boström et al., 2016; Potier et al., 2020). Whilst animals receive a massive amount of visual information, their neural computing resources are limited (Itti, 2000; Koch et al., 2006). An attention mechanism consisting of a bottom-up component (stimulus-driven, also called saliency-driven) and a top-down component (driven by voluntary goals) is thus widely used to preferentially process stimuli that are the most important for survival. For most vertebrates, searching for food and detecting threats are the primary skills needed for survival. Such rare stimuli can elicit an orienting response by the visual-motor function, which is carried out by the gaze control system (Gutfreund, 2012). In addition, this process is driven by bottom-up visual saliency, even in sedated animals (Knudsen and Schwarz, 2017a).

The saliency of a particular object results from how different it is from surrounding objects (spatial saliency mapping) (Itti et al., 1998). A large body of evidence leaves no doubt that the GABAergic isthmi pars magnocellularis (Imc) is indispensable for competitive interactions between spatial stimuli and selective attention (Mahajan and Mysore, 2019; Mysore and Knudsen, 2013; Schryver et al., 2020). Imc neurons with a stronger stimulus in their receptive field (RF) will inhibit neurons with a weaker stimulus; this inhibition is not only unaffected by the distance between the competitive neurons but is also independent of the value of the visual feature (Mahajan and Mysore, 2019; Schryver and Mysore, 2019; Schryver et al., 2020). The firing rates of Imc neurons could thus represent the relative saliency of the corresponding spatial visual stimuli, with the most salient location capturing the animal’s attention and gaining access to working memory. This process, which computes the saliency of objects in the whole visual field, is usually called spatial saliency mapping. Years of evidences show that, in birds, the midbrain selective attention network (comprising the optic tectum and nucleus isthmi) and, especially, the Imc play a key role in spatial saliency mapping, which computes the most spatially salient stimulus for the appropriate brain regions to control the body.

Saliency not only results from spatial surround competition, but also depends on how a current stimulus differs from past stimuli (Dutta and Gutfreund, 2014; Dutta et al., 2016; Garcia-Diaz et al., 2012; Gutfreund, 2012; Lee et al., 2020; Weber et al., 2019). The strength of a neuron’s response to the first presentation of a novel stimulus is much greater than to all subsequent presentations of the same stimulus. This response habituation, which is specific for the location and physical properties of the stimulus, is referred to as stimulus-specific adaptation (SSA) (Netser et al., 2011). This adaptation can facilitate the detection of out of the ordinary stimuli in the temporal domain in a process called temporal saliency mapping (Dutta and Gutfreund, 2014; Ferger et al., 2018; Gutfreund, 2012; Reches and Gutfreund, 2008; Zhai et al., 2019), which is equally important for an animal’s survival. The Imc is known to be the main participant in spatial saliency mapping and, scientific interest in its possible role in temporal saliency mapping is increasing. A previous study in pigeons has shown that the optic tectum displays visual SSA to direction of motion (Wasmuht et al., 2017). Considering the close interaction between the optic tectum and the Imc in spatial saliency mapping, important questions that remains to be addressed are whether visual SSA also exists in the Imc and if temporal saliency for direction of motion can result from that visual SSA.

The aim of this study was to clarify the role of the Imc in visual SSA and temporal saliency for direction of motion in birds. Based on previous research, we propose three possible hypotheses that can be tested: (1) visual SSA exists in the Imc in pigeons; (2) motion direction, speed of visual stimulus and inter-stimulus interval (ISI) will modulate the degree of SSA; and (3) temporal saliency for motion direction may result from visual SSA in the Imc. In the following experiments, we used a classical constant order paradigm, consisting of objects moving in opposite directions across the center of the Imc’s RF, to test these hypotheses.

## Materials and methods

### Animal preparation

Neuronal recordings were carried out in male and female pigeons (*Columba livia*, 350–450 g body weight), which were housed in individual wire mesh cages under a 12:12 h light-dark cycle with free access to water and food. All experiments were conducted in accordance with the Animals Act, 2006 (China) for the care and use of laboratory animals and were approved by the Life Science Ethical Review Committee of Zhengzhou University.

### Surgery and recording

Experiments were performed following protocols that have been described previously (Wang et al., 2020). Briefly, surgery started after the animals were anesthetized with 20% urethane (1 mL/100 g body weight). When the birds closed their eyes and no longer responded to auditory or painful stimulation, they were transferred to a stereotaxic device. Their heads were placed in the stereotaxic holder, the right eye was kept open and moisturized with saline solution during the experiment, while the left eye was covered. A small hole was then drilled in the bone to expose the left dorsal brain, allowing access to the Imc. A small slit was then made in the dura using a syringe needle, permitting dorsoventral penetration through the Imc. Throughout the experiment, a heating panel was used to maintain the pigeon’s body temperature at about 41°C.

Multi-unit activity was recorded under urethane anesthesia using 16-channel microelectrode arrays (4 × 4, Clunbury Scientific, Bloomfield Hills, Michigan, USA), which were inserted into the Imc using a micromanipulator. We targeted the Imc following previously described methods; dorsoventral penetrations through the Imc were made at a medial-leading angle of 5 degrees from the vertical to avoid a major blood vessel in the path to the Imc, and Imc targeting was validated at the outset of this study through anatomical lesions (Mahajan and Mysore, 2018).

Spike signals of units were recorded with a sampling frequency of 30 kHz and extracted with a band-pass filter (250–5 kHz). Local field potential signals were amplified (4000×), filtered (0–250 Hz) and continuously sampled at 2 kHz using a Cerebus^®^ recording system (Blackrock Microsystems, Salt Lake City, UT, USA). All data recorded were analyzed off-line using customized MATLAB applications.

### Visual stimuli

Visual stimuli were generated using the MATLAB-based Psychophysics Toolbox (Psychtoolbox-3; www.psychtoolbox.org), running on Windows, and were synchronized with the recording system. A LED monitor (112 degrees vertical × 80 degrees horizontal, running at 100 Hz) was placed tangential to, and 40 cm from, the pigeon’s right eye to present monocular visual stimuli. The luminance of the gray screen background was 118 cd/m^2^ and that of the black stimuli was 0.2 cd/m^2^.

Three visual stimuli were used for data collection. The first was a black dot (1.3 degrees) against a gray background, moving randomly at 20–40 degrees/s along a series of parallel paths over the tangent LED monitor in front of the pigeon to map the RF of Imc neurons. Before the formal experiments started, the RFs of Imc neurons were roughly estimated by manually sliding a dot delivered by a simple program around the screen, while simultaneously monitoring the firing responses. Taking into account the large RF of Imc neurons, and to maximize the use of the monitor area, we then adjusted the position and angle of the monitor to ensure that its long axis was parallel with the long axis of the RF of the Imc.

The second visual stimulus, a black dot of 1.3 degrees moving in 16 directions (temporal-nasal direction set as 0 degrees) against a gray background, spaced by 22.5 degrees, was used to measure the directional tuning curves of Imc neurons before and after the constant order paradigm. Within each trial, all the motion directions were repeated five times, following different pseudorandom orders, and the distances of motion in each direction were equal and not shorter than the long axis of the Imc RF, fitted by a two-dimensional Gaussian model.

The third visual stimulus, the constant order paradigm, was used to verify the existence of SSA in the Imc. The motion path in this test was parallel to and longer than the long axis of the Imc RF. One black 1.3 degree dot against a gray background, occurring outside the RF of Imc neurons, thus immediately started to move in one direction 20 times, followed by 20 sweeps in the opposite direction. Each 20-trial set of dots moving in the same direction was termed a block, and each block was repeated 11 times, so that the total number of trials was 440 (220 for each direction). Dots moving at 30 degrees/s and 60 degrees/s along the same path were used in the experiment, and each motion sweep was followed by an ISI of 0, 1 or 3 s at each speed.

### Histology

As described in our previous publication (Wang et al., 2020), once data recording was complete, the actual site of the recorded unit in the brain was identified using anatomical lesions. Briefly, electrolytic lesions were made to mark the site of the recorded Imc neuron by passing a 0.2 mA current through the same recording electrode for 30 s. The bird was then euthanized with 20% urethane and perfused transcardially with phosphate buffered saline followed by 4% paraformaldehyde. The brain was removed from the skull and tissues were postfixed for 12 h, followed by immersion in 30% sucrose solution at 4°C for 24 h. Finally, the brain was frozen and cut into 40-μm transverse sections on a freezing microtome and stained with Cresyl violet for subsequent microscopic observations of marked and lesioned sites.

### Analysis

The ‘Wave_clus’ spike-sorting toolbox (Quiroga et al., 2004), a fast and unsupervised algorithm for spike detection and sorting, was used to sort the recorded data into single units. We included in the analysis only those units that had less than 5% of the spikes within 1.5 ms of each other (Mahajan and Mysore, 2018). The RFs of Imc units were estimated by calculating the firing rate on the screen for each unit. The analysis window was from 80 ms after the onset of the stimulus to 80 ms after the offset of the stimulus, and the baseline response was estimated in the 250 ms before the stimulus. Neural responses were quantified as firing rate in the analysis window minus firing rate of baseline activity, and were calculated by averaging the response of the units across all trials; areas with above baseline response were selected as the RFs of the recorded units. The RF centers of the recorded units were adequately fitted by a two-dimensional Gaussian model.

In each case, for analysis of visual SSA, the first two blocks were not included in the analysis to avoid onset effects. The first presentation of a stimulus within a block was regarded as the deviant stimulus, while the last stimulus within a block was used to assess the response to the common stimulus.

To quantify the SSA effect for each direction, we used the index defined by previous researchers (Ulanovsky et al., 2003). The stimulus index (SI) for direction *i* is the normalized difference between the mean response to the deviant stimulus and the mean response to the common stimulus, defined as:

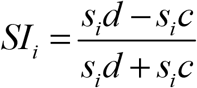

Where *s*_*i*_ *d* denotes average responses to direction *i* deviant stimuli and *s*_*i*_*c* denotes average responses to direction *i* common stimuli. In the following description, the stimulus indices for directions 1 and 2 are abbreviated SI1 and SI2, respectively. As pointed out in previous literature (Ulanovsky et al., 2003), if the adaptation is not stimulus-specific, the sum of SI1 and SI2 is expected to be zero. Thus, units for which the sum of SI1 and SI2 is greater than zero is a widely accepted criterion for SSA at the population level (Hershenhoren et al., 2014; Malmierca et al., 2009; Taaseh et al., 2011; Valdes-Baizabal et al., 2017). In addition, recorded units for which both SI1 and SI2 are greater than zero show stronger responses to the first deviant for both directions (Wasmuht et al., 2017). To analyze the population tendency for SSA, population averages were obtained by normalizing each recorded site by the maximum response and computing the mean across recorded sites.

To explore the time course of visual SSA in Imc neurons that have the same stimulus duration, we generated post-stimulus time histograms (PSTHs), with 20 ms bins for the first and last presentation of the same directional motion in the sequence. PSTHs were normalized to the maximum bin and averaged across the population. PSTHs were smoothed for display purposes (running average of five adjacent points).

Since previous studies have suggested that recent stimuli may alter directional tuning (Vinken et al., 2017), we tested the tuning curve before and after each complete trial. We thus introduced the concept of change ratio (CR) of response strength to quantify the effect of repetitive directional moving dots on the directional tuning curves, defined as:

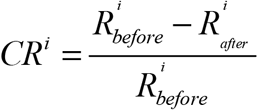

where 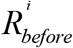 denotes the response strength for direction *i* before the constant paradigm and 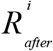 represents the response strength for that direction after the constant paradigm.

To test the significance of habituation, we performed t tests to compare the population average response of the first stimulus in the sequence with the average responses to all the remaining stimuli. A Holm–Bonferroni multiple-comparison correction was applied to all tests. To test the speed dependence of visual SSA, we used the *ANOVA* test to compare population average sums of SI1 and SI2 under different speeds and we used the trend test (Cleophas and Zwinderman, 2016) to describe the parameter dependence of the degree of visual SSA in the Imc.

## Results

Data presented here were collected from Imc single- and multi-units in fifteen anesthetized pigeons. Both spike and local field potential signals of all units were stored simultaneously throughout the experiments. The multi-units data were then subjected to spike sorting (see Materials and methods) for all remaining analyses in this study.

### Receptive field mapping and directional tuning of example Imc unit

Since previous researches have shown that Imc neurons have elongated RFs (Li et al., 2007; Wang and Frost, 1991), for which the ratio of the long axis to the short axis can exceed three, only units with a typical RF were included in the following analyses. The RF of a sample unit covered almost 100 degrees in height and 30 degrees in width (Figure 1A). The directional tuning curves of the recorded neurons in this study were measured before and after the constant order paradigm. Imc neurons that we recorded mostly responded strongly to upward (v-d) and downward (d-v) motion, with a weak response in the forward (t-n) direction and very little, if any, response in the backward (n-t) direction (Figure 1B). These results are in good agreement with previous research (Li et al., 2007; Mysore and Knudsen, 2013; Wang and Frost, 1991) and, given that a dot moving upwards and downwards can evoke a robust response, we took the upward direction as direction 1 and the downward direction as direction 2 for subsequent constant order paradigms. After completing the experimental procedure, the actual position of the recorded unit in the Imc was verified as shown in Figure 1C.

**Figure 1.**
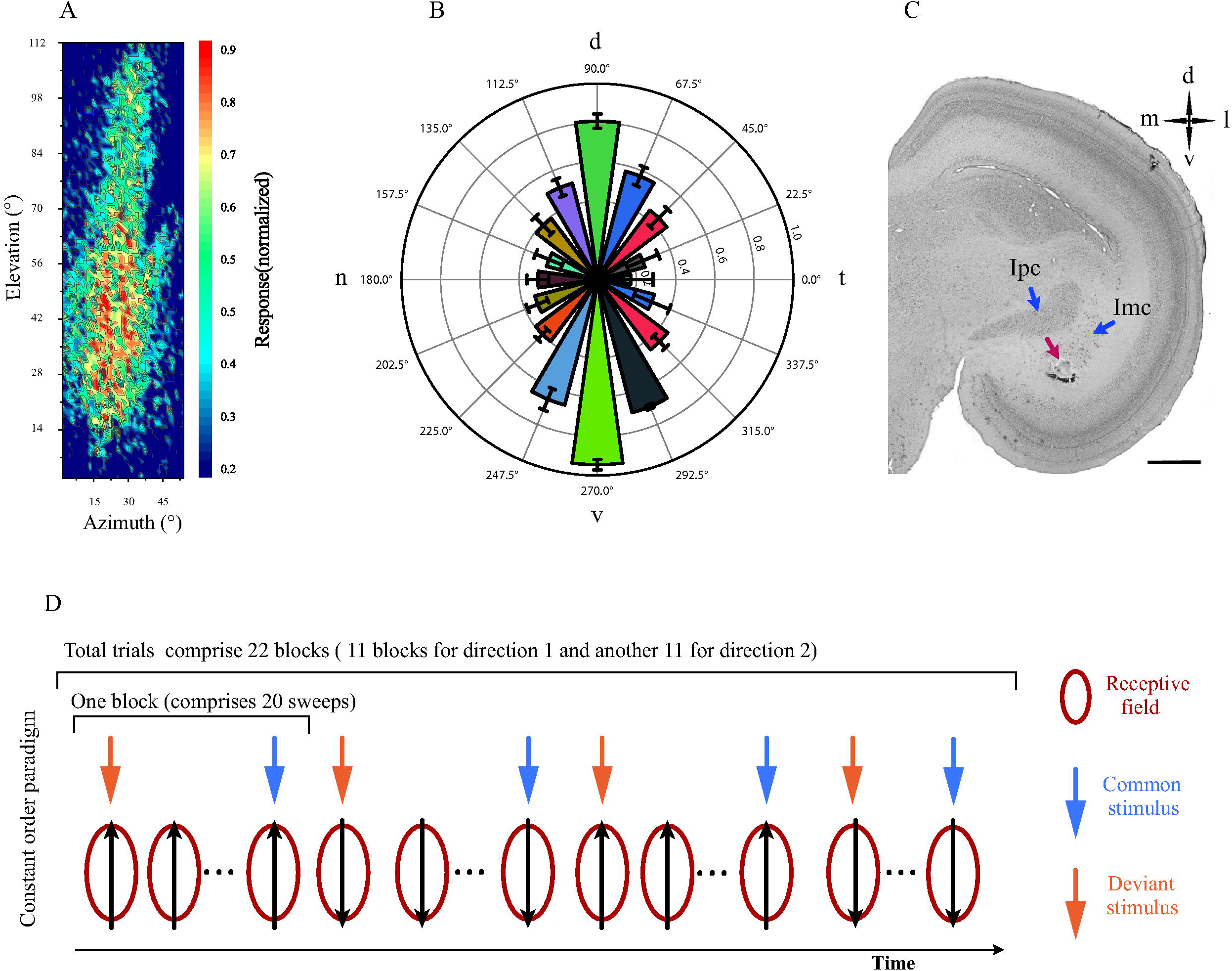
Response Properties of example Imc unit and constant order experimental paradigm. (A) Strengths of response of example Imc unit in spatial receptive field, normalized to maximum response (range approximately 30 degrees horizontally and 100 degrees vertically); (B) Directional tuning of example unit when the monitor was rotated through 38 degrees to meet the normal angle between the horizontal meridian and the pigeon’s bill. The stimulus was a black dot, presented on a gray background, moving at 30 degrees/s in 16 directions, spaced by 22.5 degrees. Each direction of motion was repeated 10 times. Error bars = standard deviation; (C) Bright field photomicrograph of 40-μm Nissl-stained coronal section, showing recording site of sample unit, Scale bar = 1000 μm. d, dorsal; l, lateral; m, medial; n, nasal; t, temporal; v, ventral; (D) Constant order paradigm used to test the existence of SSA in this study. Upward and downward motions were selected to alternately cross the receptive field (red circle) of the Imc unit. Each block consists of 20 sweeps in the same direction (shown as black arrows across the red circles), followed by a block of 20 sweeps in the opposite direction. The total trials comprised 11 alternations. The path of the upward direction overlapped with that of the downward direction. The first presentation of one direction of motion in each block (indicated by orange arrows) was set as the deviant stimulus, and the twentieth in that block was regarded as the common stimulus (indicated by blue arrows).

### Imc shows long lasting adaptation to visual motion direction

Adaptation has been shown to exist widely in many vertebrate sensory systems, including the auditory system (Parto Dezfouli et al., 2019; Reches and Gutfreund, 2008; Zhai et al., 2019), and recent research has provided evidence that visual adaptation also exists in the optic tectum of birds (Vinken et al., 2017; Wasmuht et al., 2017). The response of an example Imc unit to a constant order paradigm (Figure 1D), consisting of a dot moving at 30 degrees/s and a 1 s ISI, is shown in Figure 2A. The firing rate increased when the dot swept through the RF in both upward and downward directions. Within the sequence, the strength of the response decreased rapidly when the same direction of motion was repeated several times. Across the trials at 30 degrees/s, the average response to the first presentation (deviant stimulus) after switching directions was significantly larger for both direction 1 and direction 2 (*t* test, p <0.05) than the average response to the last stimuli of each block (Figure 2B), implying some degree of adaptation. The response of the example Imc neuron to the first presentation of direction 1 was significantly greater than that to the last presentation of direction 1 (Figure 2C, t test, p <0.05).

**Figure 2.**
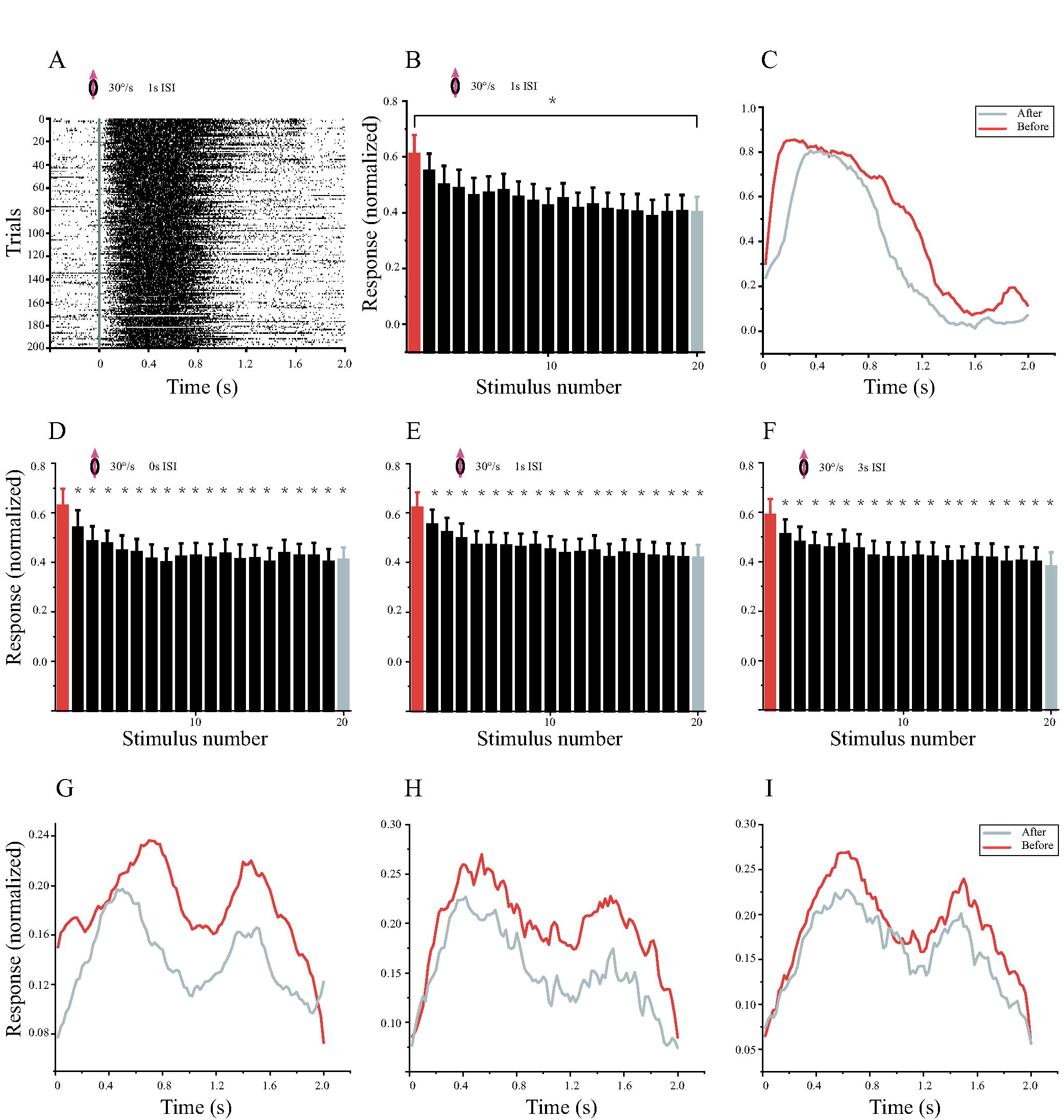
Visual adaptation of Imc units to direction of motion. (A) Spike raster plots of example Imc unit to motion in upward direction. Each row represents plot of one sweep and plots are arranged as a function of the order of the stimulus within total blocks. The vertical gray line indicates the onset of each sweep; (B) Histogram of example unit’s normalized average responses as a function of the order of the stimulus within the sequence. Error bars show 95% confidence interval. The asterisk indicates that response to the first stimulus was significantly greater than that to the last stimulus (t test, p <0.05); (C) PSTH of example unit’s response to the first stimulus (orange curve) compared with PSTH of response to the last stimulus in the sequence (38 s later, gray curve); (D) Population response of neurons to upward motion at 30 degrees/s, with stimulation duration of 2 s and ISI of 0 s. Histogram shows normalized population response to stimulation as a function of position within the block. Orange column in the histogram represents responses to the first upward motion and gray column represents response to the last upward motion. Asterisk indicates response that was significantly smaller than response to the first stimulus (t test, Holm–Bonferroni correction; p <0.05). Error bars show 95% confidence interval; (E) Population response of neurons to upward motion at 30 degrees/s with stimulation duration of 2 s and ISI of 1 s. Histogram shows normalized population response to stimulation as a function of position within the block. Format same as in D; (F) Population response of neurons to upward motion at 30 degrees/s with stimulation duration of 2 s and ISI of 3 s. Histogram shows normalized population response to stimulation as a function of position within the block. Format same as in D; (G) Population PSTH of response to the first stimulus (orange curve) significantly surpassed (t test, p <0.05) population PSTH of response to the last stimulus in the sequence (38 s later, gray curve) for D; (H) Population PSTH of response to the first stimulus (orange curve) significantly surpassed (t test, p <0.05) PSTH of response to the last stimulus in the sequence (57 s later, gray curve) for E; (I) Population PSTH of response to the first stimulus (orange curve) significantly surpassed (t test, p <0.05) population PSTH of response to the last stimulus in the sequence (95 s later, gray curve) for F.

Previous researches have shown multiple time scales of adaptation and a long lasting specific adaptation (Netser et al., 2011; Ulanovsky et al., 2004). To explore the time course of visual SSA in the Imc, we selected units with the same stimulus distance for population analysis. As shown in Figure 2D, the response to the first presentation of motion in direction 1 at 30 degrees/s and 0 s ISI was significantly greater than that to following stimuli (*t* test with Holm–Bonferroni correction, p <0.05). A plot of population PSTH of the response to the first sweep against that of the response to the last sweep, 38 s later (Figure 2G) showed a marked change in response strength (*t* test, p <0.05). For the 1 s and 3 s ISIs, the response to the first presentation of motion in direction 1 at 30 degrees/s was also significantly greater than the response to subsequent stimuli (Figure 2E and Figure 2F, *t* test with Holm–Bonferroni correction, p <0.05). The response strength to the first presentation differed significantly from the last sweep with 1 s and 3 s ISIs (Figure 2H and Figure 2I, 57 s and 95 s later, respectively, *t* test, p <0.05). These results show that a preceding sweep of up to 95 s was sufficient to significantly reduce the response to the same motion direction, indicating a strong and long-lasting neural adaptation in Imc neurons.

### Specificity of long-lasting adaptation in Imc to visual motion direction

Considering the close interaction between the optic tectum and the nucleus isthmi (especially the Imc), it is natural to assume the existence of SSA in the Imc. To confirm if this long lasting adaptation to motion at 30 degrees/s with 1 s ISI is stimulus-specific, we used a classical paradigm to test whether Imc neurons would show visual SSA to motion direction. As shown in Figure 3A, a strong response was observed to the first presentation (deviant stimulus) of one motion direction in each block, and the response then decreased on repeated trials. SIs for directions 1 and 2 were calculated to be 0.17 and 0.08, respectively. This adaptation was thus stimulus-specific according to a widely accepted criterion (Ulanovsky et al., 2003) for SSA, which requires the sum of SI1 and SI2 to be greater than zero. Different speed/ISI pairings were used to rule out the possibility of serious specific parameter dependence of the visual SSA. The average response of a sample neuron responding to motion at 60 degrees/s and 1 s ISI is shown as a histogram in Figure 3B. The strength of the response to the first presentation was significantly larger for both direction 1 and direction 2 than the last one in each block (*t* test, p <0.05). The SIs for direction 1 and direction 2 were 0.44 and 0.13, respectively, showing that adaptation here was also stimulus-specific.

**Figure 3.**
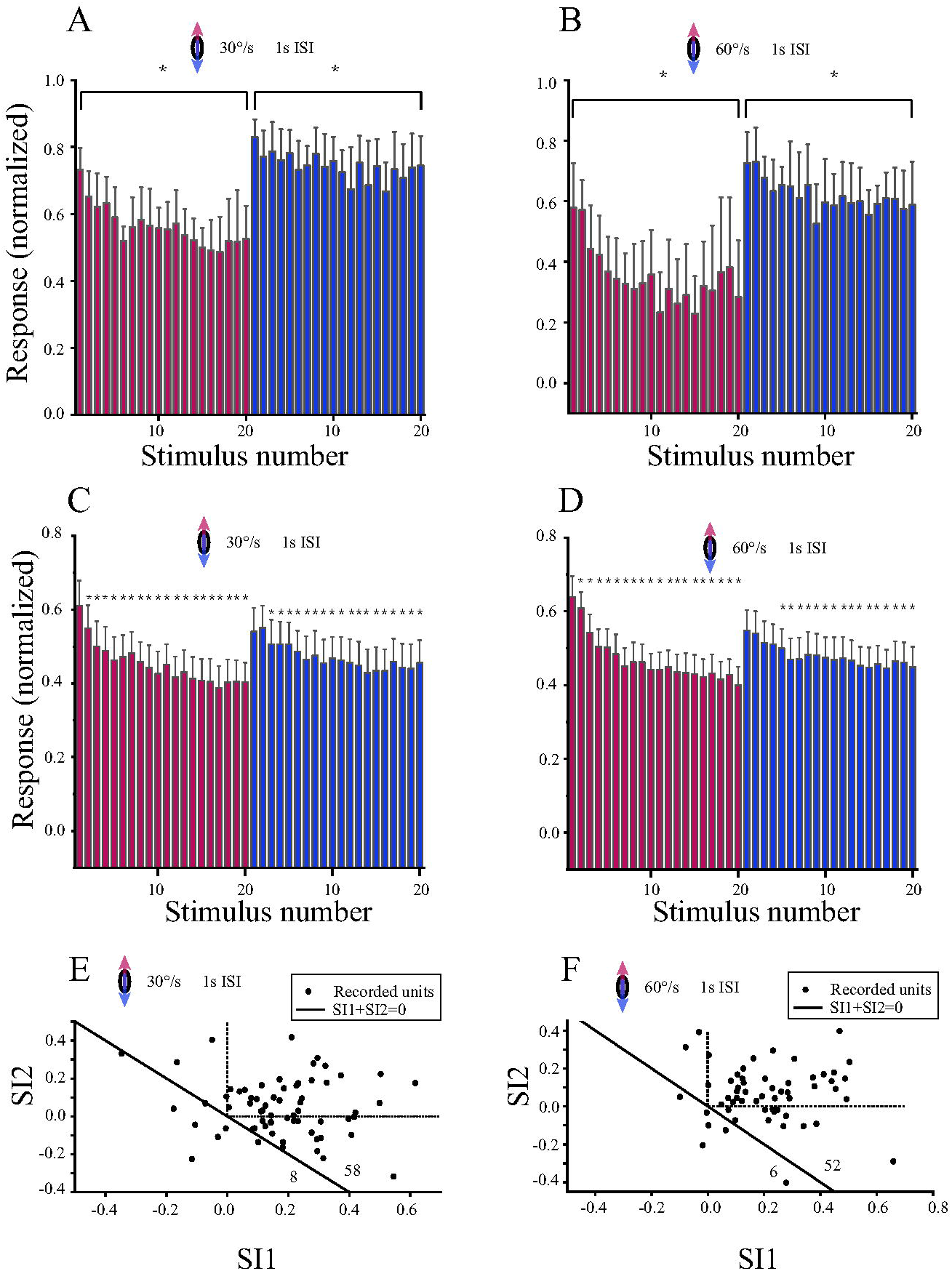
Visual SSA to direction of motion observed in Imc units. (A) Visual SSA occurs in the constant order paradigm at 30 degrees/s, with a 1 s ISI. Histogram shows normalized average response of example Imc unit (in Figure 2) as a function of the order of the stimulus within the sequence. Red columns represent upward motion and blue columns represent downward motion, with overlapping paths. Asterisks indicate that responses to the first stimulus were significantly greater than to the last stimulus (t test, p <0.05). Error bar shows 95% confidence interval; (B) Visual SSA occurs in the constant order paradigm at 60 degrees/s, with a 1 s ISI. Format same as in A; (C) Population average responses to visual upward and downward motion at 30 degrees/s with a 1 s ISI. Histogram shows normalized average responses as a function of the order of the stimulus within the sequence. Orange columns represent responses to upward motions and blue columns represent responses to downward motions. Error bar shows 95% confidence interval. Asterisks indicate responses that were significantly smaller than the response to the first stimulus (t test, Holm–Bonferroni correction; p <0.05). Red columns represent upward motion and blue columns represent downward motion, with overlapping paths; (D) Population average responses to visual upward and downward motion at 60 degrees/s with a 1 s ISI. Format same as in C; (E) Scattergram showing SIs for motion at 30 degrees/s with a 1 s ISI in direction 1 plotted against SIs for motion at 30 degrees/s with a 1 s ISI in direction 2. The diagonal solid line shows that the sum of SI1 and SI2 is equal to zero. The dashed lines represent the upper right quadrant where SIs for both directions are positive. The numbers at the lower right corner indicate the number of points above and below the diagonal line; (F) Scattergram showing SIs for motion at 60 degrees/s with a 1 s ISI in direction 1 plotted against SIs for motion at 60 degrees/s with a 1 s ISI in direction 2. Format same as in E. Asterisks represent responses that were significantly smaller than the response to the first stimulus (t test, Holm–Bonferroni correction; p <0.05).

To investigate visual SSA at the population level, response data collected at 30 degrees/s and 60 degrees/s (both with a 1 s ISI) were averaged across trials and units. As shown in Figure 3C, the strength of the average response to the first appearance of directional motion was significantly larger than the average response strength to most subsequent stimuli for either direction (t test with Holm–Bonferroni correction, p<0.05). Similar results were observed in the population average response to motion at 60 degrees/s with a 1 s ISI (Figure 3D). The average response to the first presentation of motion in one direction was significantly greater than those to most subsequent stimuli in either direction (t test with Holm–Bonferroni correction, p <0.05). SIs for both conditions were calculated, and SI2 was plotted against SI1 for each unit (Figure 2E for recorded units in Figure 2C, Figure 2F for recorded units in Figure 2D). The scatter plot in Figure 2E shows that 87% of the recorded units lay above the diagonal line, indicating SSA to motion direction at 30 degrees/s with a 1 s ISI. The proportion for motion at 60 degrees/s and a 1 s ISI was 90%, implying visual specific adaptation to direction of motion and confirming our first hypothesis.

### Inter-stimulus interval, direction and speed dependence of visual SSA in Imc

To investigate dependence of visual SSA on the ISI, speed and direction, we used a dot moving along the same path in both directions, with other ISI/speed pairings. Plots of population average responses for a dot moving at 60 degrees/s with a 0 s ISI and at 60 degrees/s with a 3 s ISI are shown in Figure 4A and Figure 4B, respectively. The response to the first presentation was significantly stronger than most subsequent stimuli in either direction (t test with Holm–Bonferroni correction, p <0.05). Scattergrams of SI2 against SI1 (insets in Figure 4A and Figure 4B) show that 91% of the recorded units show visual SSA to motion at 60 degrees/s with a 0 s ISI and that 81% show visual SSA with a 3 s ISI. There was also a significant decreasing trend for population sums of SI1 and SI2 at 60 degrees/s (Figure 4C) with increasing ISIs (p trend <0.05), showing the dependence of visual SSA in the Imc on ISI.

**Figure 4.**
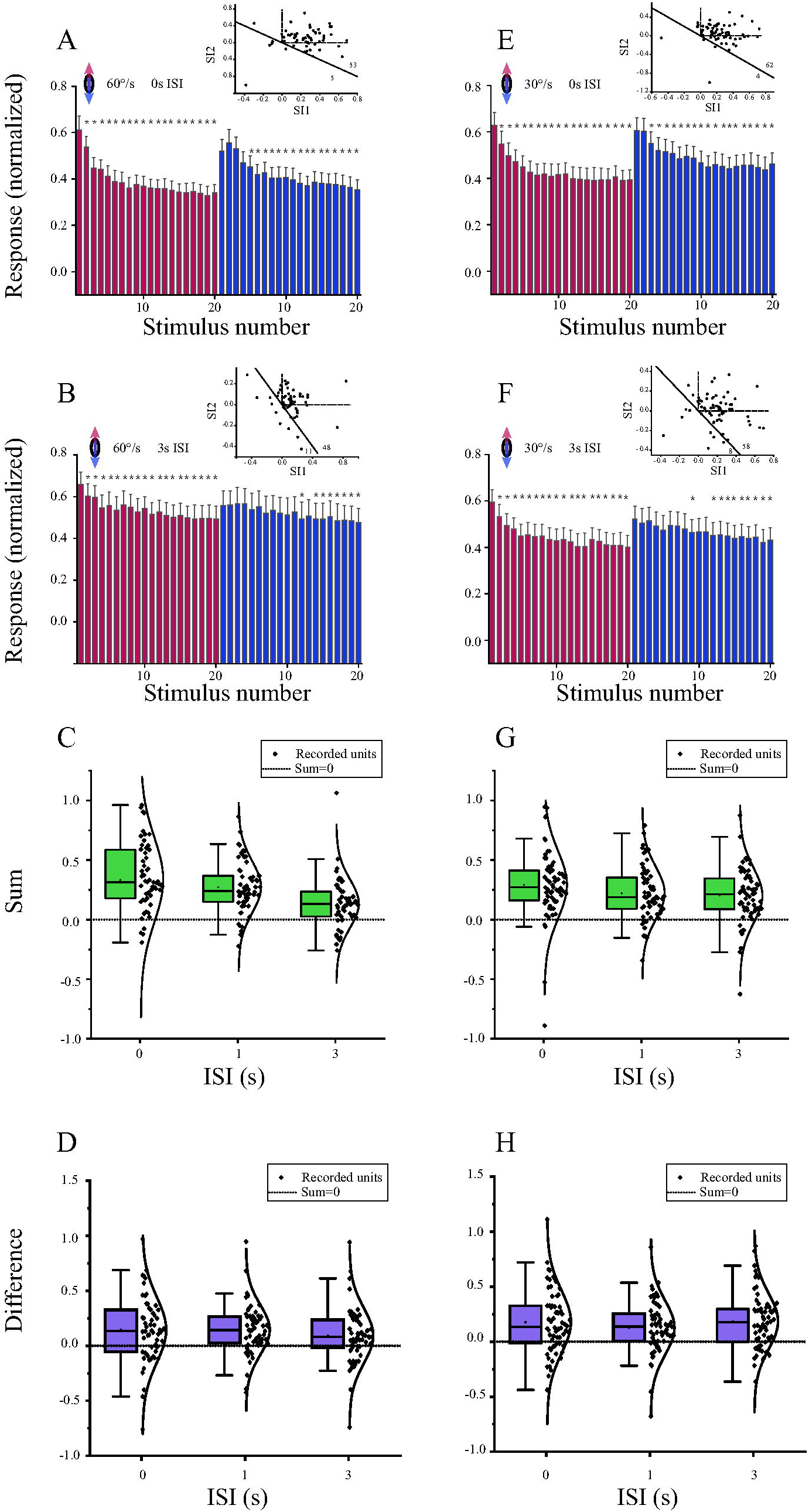
Dependence of degree of SSA on direction, speed and inter-stimulus interval. (A) Average responses to upward and downward motion at 60 degrees/s and a 0 s ISI along same path. Histogram shows responses sorted by trial order within the block. Red columns represent upward motion and blue columns represent downward motion, with overlapping paths. Scattergram in inset shows SIs for upward direction plotted against SIs for downward direction. The diagonal solid line shows that the sum of SI1 and SI2 is equal to zero. The dashed lines represent the upper right quadrant where SIs for both directions are positive. The numbers at the lower right corner indicate the number of points above and below the diagonal line. Error bar shows 95% confidence interval. Asterisks indicate responses that were significantly smaller than the response to the first stimulus (t test, Holm–Bonferroni correction; p <0.05); (B) Average responses to upward and downward motion at 60 degrees/s with a 3 s ISI along the same path. Histogram shows responses sorted as a function of the order of the stimulus within the sequence. Format same as in A; (C) Boxplot showing sums of SI1 and SI2 to motion at 60 degrees/s plotted against ISI. Values are significantly greater than zero (t test, p <0.05), in agreement with the linear trend (trend test, p <0.05). Whisker indicates 1.5 times the interquartile range; (D) Boxplot showing values of SI1 minus SI2 to motion at 60 degrees/s for each unit plotted against ISI. Values are significantly greater than zero (t test, p <0.05). Format same as in C; (E) Average responses to upward and downward motion at 30 degrees/s with a 0 s ISI along same path. Histogram shows responses sorted by trial order within the block. Format same as in A; (F) Average responses to upward and downward motion at 30 degrees/s with a 3 s ISI along same path. Histogram shows responses sorted as a function of the order of the stimulus within the block. Format same as in A; (G) Boxplot showing sums of SI1 and SI2 to motion at 30 degrees/s plotted against ISI. Values are significantly greater than zero (t test, p <0.05), in weak agreement with linear trend (trend test, p = 0.055). Format same as in C; (H) Boxplot showing values of SI1 minus SI2 to motion at 30 degrees/s for each unit plotted against ISI. Values are significantly greater than zero (t test, p <0.05). Format same as in D.

SI2 was greater than SI1 for the example Imc unit (Figures 2B and 2C), implying dependence of visual SSA on direction. To examine this trend at the population level, we calculated the difference between SI1 and SI2 for each recorded unit. The population difference was significantly (*t* test, p <0.05) greater than zero for all ISIs at 60 degrees/s (Fig. 4D), in agreement with the result obtained with the sample neuron, implying direction dependence of the degree of visual SSA in the Imc.

For motion at 30 degrees/s (Figure 4E with a 0 s ISI and Figure 4F with a 3 s ISI), the population response to the first occurrence of motion direction was significantly greater than almost all the subsequent stimuli for both ISIs and directions (t test with Holm–Bonferroni correction, p <0.05). As the insets in Figures 4E and 4F show, 94% of the recorded units showed visual SSA to motion at 30 degrees/s with a 0 s ISI, and that proportion was 88% with a 3 s ISI. There was also a weak decreasing trend for population sums of SI1 and SI2 at 30 degrees/s (Figure 4G) with increasing ISIs (p trend = 0.055). Similar results were observed for motion at 30 degrees/s; SI1 was significantly greater than SI2 at the population level for all ISIs (Figure 4H, *t* test, p <0.05), showing that visual SSA in the Imc is dependent on direction. Lastly, to investigate the speed dependence of visual SSA in the Imc, we compared population distributions of the sum of SI1 and SI2 to motion at 30 degrees/s and 60 degrees/s with the same ISIs (Figures 4E and 4F). Weak differences were observed between the two speeds for a 0 s ISI (one-way ANOVA, p = 0.47), a 1 s ISI (one-way ANOVA, p = 0.19) and a 3 s ISI (one-way ANOVA, p = 0.08).

### Temporal saliency for motion direction resulting from SSA in Imc

Previous studies have shown that SSA in other sensory systems leads to a shift in directional tuning (Ogawa and Oka, 2015; Shen et al., 2015). For the example unit (Figure 5), we thus also measured directional tuning curves after the constant order paradigm. We observed marked center decrements and surround increments with regard to direction 1 (v-d) and direction 2 (d-v) for responses before the constant order paradigm at both speeds (Figures 5A and 5E). To explore the distributions of response changes in the neural population, we tested the directional tuning curve of each recorded unit before and after the constant order paradigm. The change ratio was calculated across the recorded Imc neurons for all parameter pairs (as described in Materials and methods). As shown in Figures 5B-D and Figures 5F-H, significant attenuation of the response (*t* test, p <0.05) to directions 1 and 2 was introduced by the constant order paradigm in all cases. The surround directions relative to directions 1 and 2, especially the 157.5 degrees direction and t-n direction (Figure 1B), also showed significant increments after the paradigm (*t* test, p <0.05). Similar results were also observed at 60 degrees/s with all ISIs. In the optic tectum of barn owls, an enhancement of rare stimuli was observed in long-lasting adaptation to an auditory stimulus but not in short-lasting adaptation (Netser et al., 2011), and this similar long-lasting SSA to repetitive visual motion direction enhanced salient detection for novel motion directions in Imc neurons.

**Figure 5.**
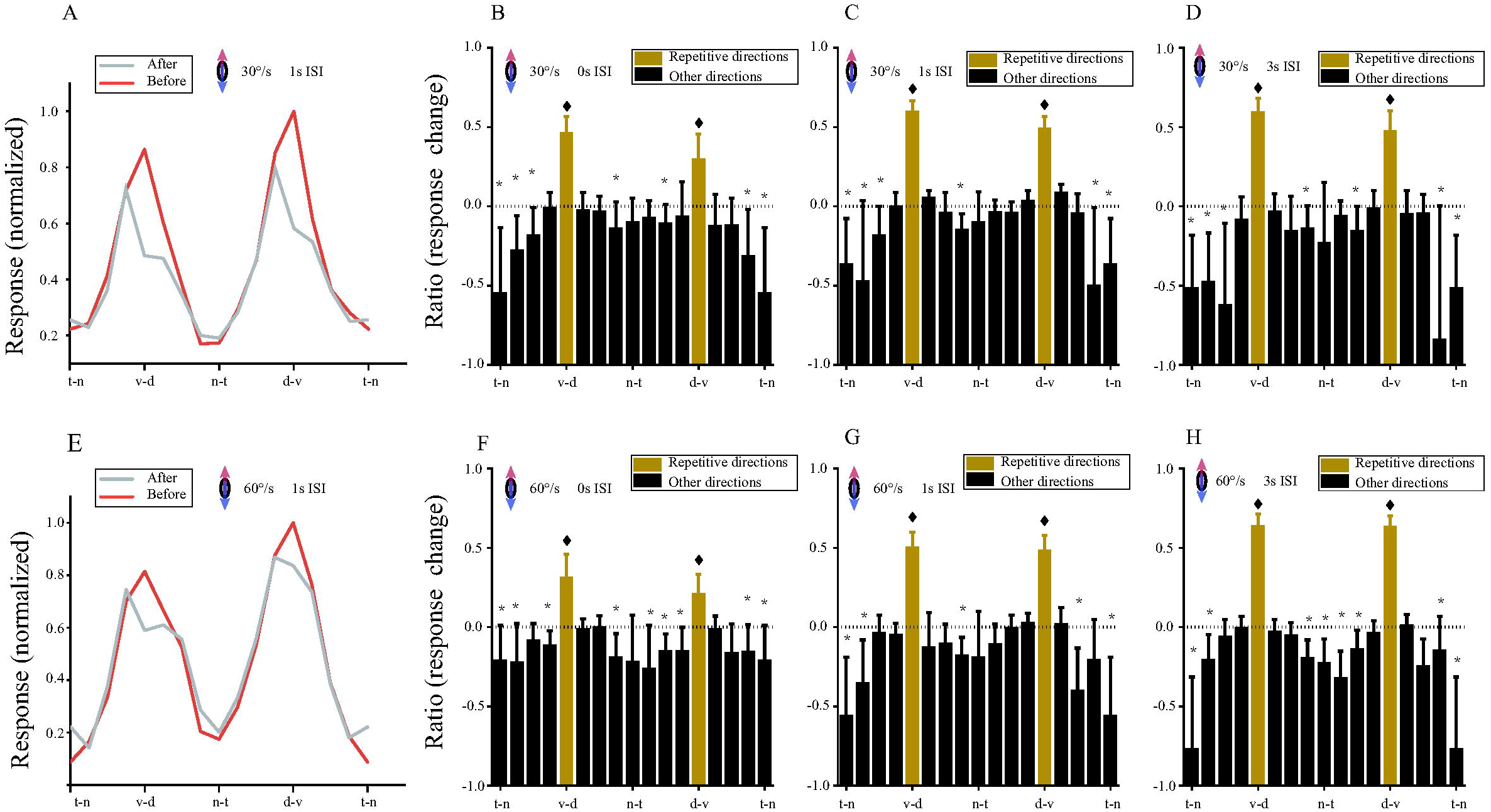
Detection of salient motion direction from SSA in Imc. (A) Directional tuning curves of example unit (in Figure 2) before and after constant order paradigm with visual motion at 30 degrees/s and a 1 s ISI. Orange line represents before visual stimuli and gray line denotes after visual stimuli; (B) Change ratios of response strength in directional tuning at population level for different directions at speed of 30 degrees/s with a 0 s ISI. The dotted line shows that the change ratio is zero. Error bar shows 95% confidence interval. Asterisks indicate change ratios that are significantly smaller than zero (t test, p <0.05). Diamonds represent change ratios that are significantly greater than zero (t test, p <0.05); (C) Change ratios of response strength in directional tuning at population level for different directions at speed of 30 degrees/s with a 1 s ISI. Format same as in B; (D) Change ratios of response strength in directional tuning at population level for different directions at speed of 30 degrees/s with a 3 s ISI. Format same format as in B; (E) Change ratios of response strength in tuning curves of example neuron (in Figure 2) before and after constant order paradigm with visual motion at 60 degrees/s and a 1 s ISI. Orange line represents before visual stimuli and gray line indicates after visual stimuli. Format same as in A; (F) Change ratios of response strength in directional tuning at population level for different directions at speed of 60 degrees/s with a 0 s ISI. The dotted line shows that the change ratio is zero. Error bar shows 95% confidence interval. Asterisks indicate change ratios that are significantly smaller than zero (t test, p <0.05). Diamonds represent change ratios that are significantly greater than zero (t test, p <0.05); (G) Change ratios in directional tuning at population level for different directions at speed of 60 degrees/s with a 1 s ISI. Format same as in (F); (H) Changes in directional tuning changes at population level for different directions at speed of 60 degrees/s with a 3 s ISI. Format same as in (F).

## Discussion

Although the existence of SSA in the auditory system of many vertebrates has been clearly demonstrated (Parto Dezfouli et al., 2019; Reches and Gutfreund, 2008; Zhai et al., 2019), there have been few reports of visual SSA (Dutta et al., 2016; Netser et al., 2011; Wasmuht et al., 2017). The Imc, which has been demonstrated to play a key role in competitive interactions between spatial stimuli and selective attention (Schryver et al., 2020), is an ideal model for exploring the neural basis of visual SSA. Whereas the focus of previous experiments was the role that the nucleus plays in spatial saliency mapping, here we have demonstrated that Imc neurons can adapt specifically to direction of motion. Visual SSA for direction of motion suggests that the response may represent the degree of novelty compared with past stimuli, rather than an absolute value of the visual feature. Temporal saliency for visual motion direction resulting from visual SSA can thus be coded in the Imc, which means that the GABAergic nucleus itself implements the computation of both visual spatial (Schryver and Mysore, 2019; Schryver et al., 2020) and temporal saliency. To the best of our knowledge, this is the first attempt to demonstrate that an inhibitory node can combine visual spatial (Schryver and Mysore, 2019; Schryver et al., 2020) and temporal saliency.

Most of the Imc neurons that we recorded adapted to repetitive motion direction by decreasing the strength of the response to a low level. The neurons can recover their response strength when a novel direction of motion occurs, showing long-lasting visual SSA for motion direction. Since previous studies showed that SSA can be sensitive to ISI and stimulus speed (Ferger et al., 2018; Hershenhoren et al., 2014; Ogawa and Oka, 2015; Wasmuht et al., 2017), we tested that point by using different ISIs, whilst keeping the direction and duration of motion the same. As shown in Figure 4, variation of ISI did not eliminate visual SSA for Imc neurons, but longer intervals reduced the degree of SSA to specific direction of motion (Figure 4). The results described above also rule out the possibility that the emergence of SSA is due to specific ISI and speed. In the population response (Figures 3 and 4), a considerable proportion of the recorded neurons in our experiments lay above the diagonal line, which is widely accepted as the criterion for SSA, and quite a few neurons (data points within the top-right quadrant of the insets) responded intensely to the first deviant in both directions. The degree of visual SSA at the population level gradually decreased when the interval increased from 0 s to 3 s at both speeds, implying that visual SSA is dependent on ISI but only weakly dependent on speed, if at all (Figures 4E and 4F).

In this constant order paradigm, upwards (direction 1) and downwards (direction 2) motion were selected to test the existence of visual SSA. As shown in Figures 4G and 4H, the population degree of visual SSA for direction 1 was significantly greater than that for direction 2. There are two possible explanations for the smaller SI for downwards motion. One possibility is that it is more difficult for Imc units to adapt their response to downwards motion, and the other possibility is that they adapt, but this adaptation is not stimulus-specific. If the latter explanation is correct, the responses to the downward deviant stimulus should not stand out from the previous responses and vice versa. Hence, a smaller SSA effect for direction 2 means that the Imc neuron adapts less easily to an object moving downwards, leading to significant direction-dependence of the visual SSA. This direction-dependence of the degree of adaptation may have an ecological significance for the survival of prey animals, since it keeps the animal vigilant at all times for predators diving from a higher visual space.

As previous researchers have demonstrated, the avian midbrain selective attention network plays a key role in controlling the animal’s gaze shift (Asadollahi and Knudsen, 2016; Goddard et al., 2014; Knudsen, 2011; Knudsen and Schwarz, 2017b; Mysore and Knudsen, 2011). Recent research suggests that visual SSA exists in the optic tectum, but it is not currently known if there is any link between visual SSA in the Imc and that in the optic tectum. Although there are differences between this study and previous research (Wasmuht et al., 2017) in terms of the number of trials within one block (20 versus 10) and selection of common stimuli (last sweep versus average of the last three sweeps in each block), we also tried to compare results from most similar parameter settings. The proportions of recorded units that showed visual SSA to directional motion at 30 degrees/s and 60 degrees/s with a 1 s ISI are very similar (87.9% and 89.7%, respectively, in the Imc versus 83.3% and 90%, respectively, in the optic tectum). Additionally, that proportion at 60 degrees/s with a 1 s ISI was greater than that at 30 degrees/s with a 1 s ISI in both the Imc and optic tectum. The difference between the Imc and the optic tectum lies in the direction dependence of the degree of visual SSA, which may result from a feedforward connection from the optic tectum to the Imc or from the inherent integration executed by local circuits in the Imc. This is another open question to be investigated in future studies.

Since recent stimulation has been shown to enhance the response to novel stimuli in other sensory systems (Ogawa and Oka, 2015; Shen et al., 2015; Vinken et al., 2017), we also included changes of response strength in other directions in our analysis, although this was neglected in the previous study in the optic tectum (Wasmuht et al., 2017). In the current study, we meticulously investigated the adaptive changes in directional tuning caused by the constant order paradigm in the midbrain inhibitory nucleus. As shown in Figure 5, the directional tuning curves of the Imc changed markedly, especially for the directions used in our constant order paradigm. Responses to motion in the v-d and d-v directions were significantly reduced after repetitive sweeps, whilst responses to motion in the t-n direction and at 157.5 degrees were markedly enhanced. The Imc’s ability to accurately adjust the response for specific directions also suggests high directional resolution, which is important for computing a novel direction of motion. Although it has proven challenging to observe both suppression and enhancement of responses using invasive action potential recordings in animals (Vinken et al., 2017), our results here supplement other successful cases and may provide a mechanism to enhance saliency detection for different directions of motion.

## Abbreviations

Imc: Isthmi pars magnocellularis
OT: Optic tectum
SSA: Stimulus-specific adaptation
ISI: Inter-stimulus interval

## Authors’ contributions

JTW: Conceptualization, Software, Investigation. SMH: Writing - original draft. ZZW: Conceptualization, Software. SWW: Data curation, Visualization. LS: Validation, Writing - review editing, Supervision. All authors read and approved the final manuscript.

## Availability of data and material

The data that support the findings of this study are available on request from the corresponding author.

## Code availability

The code that support the findings of this study are also available on request from the corresponding author.

